# Analysis of >30,000 abstracts suggests higher false discovery rates for oncology journals, especially those with low impact factors

**DOI:** 10.1101/500660

**Authors:** LM Hall, AE Hendricks

## Abstract

**Background:** Recently, there has been increasing concern about the replicability, or lack thereof, of published research. An especially high rate of false discoveries has been reported in some areas motivating the creation of resource-intensive collaborations to estimate the replication rate of published research by repeating a large number of studies. The substantial amount of resources required by these replication projects limits the number of studies that can be repeated and consequently the generalizability of the findings.

**Methods and findings:** In 2013, Jager and Leek developed a method to estimate the empirical false discovery rate from journal abstracts and applied their method to five high profile journals. Here, we use the relative efficiency of Jager and Leek’s method to gather p-values from over 30,000 abstracts and to subsequently estimate the false discovery rate for 94 journals over a five-year time span. We model the empirical false discovery rate by journal subject area (cancer or general medicine), impact factor, and Open Access status. We find that the empirical false discovery rate is higher for cancer vs. general medicine journals (p = 5.14E-6). Within cancer journals, we find that this relationship is further modified by journal impact factor where a lower journal impact factor is associated with a higher empirical false discovery rate (p = 0.012, 95% CI: −0.010, −0.001). We find no significant differences, on average, in the false discovery rate for Open Access vs closed access journals (p = 0.256, 95% CI: −0.014, 0.051).

**Conclusions:** We find evidence of a higher false discovery rate in cancer journals compared to general medicine journals, especially for those with a lower journal impact factor. For cancer journals, a lower journal impact factor of one point is associated with a 0.006 increase in the empirical false discovery rate, on average. For a false discovery rate of 0.05, this would result in over a 10% increase to 0.056. Conversely, we find no significant evidence of a higher false discovery rate, on average, for Open Access vs. closed access journals from InCites. Our results identify areas of research that may need additional scrutiny and support to facilitate replicable science. Given our publicly available R code and data, others can complete a broad assessment of the empirical false discovery rate across other subject areas and characteristics of published research.

## Introduction

Increasing concern about the lack of reproducibility and replicability of published research (1–8) has led to numerous guidelines and recommendations including the formation of the National Academies of Sciences, Engineering, and Medicine committee (9) on Reproducibility and Replicability in Science (10–13). In addition, efforts have been made to estimate the replication rate by forming large-scale collaborations to repeat a set of published studies within a particular discipline such as psychology (6), cancer biology (14), economics (15), and social sciences (16, 17). The proportion of studies that replicate vary from approximately 1/3 to 2/3 depending, in part, on the power of the replication studies, the criteria used to define replication, and the proportion of true discoveries in the original set of studies (18).

These replication projects are often massive undertakings necessitating a large amount of resources and scientists. The sheer amount of resources needed can become a barrier limiting both the number and breadth of studies repeated. Indeed, the Cancer-Biology Reproducibility project lowered its projected number of studies for replication from 50 to 37 and recently lowered the number again to 18 (19). This suggests that an efficient, complementary approach to evaluate replicability would be highly beneficial.

The false discovery rate, which is the number of scientific discoveries that are false out of all scientific discoveries reported, is a complementary measure to replicability as we expect a subset of true discoveries to replicate, but do not expect false discoveries to replicate. In 2013, Jager and Leek (20) published a method to estimate the empirical false discovery rate of individual journals using p-values from abstracts. Compared to the resource intensive replication studies mentioned above, Jager and Leek’s method is quite efficient. Here, we take advantage of this efficiency to gather and use p-values from over 30,000 abstracts to estimate the empirical false discovery rate for over 90 journals between 2011-2015. Using these journals, we evaluate if and how the empirical false discovery rate varies by three journal characteristics: (1) subject area – cancer vs. general medicine; (2) two-year journal impact factor (JIF), and (3) Open Access vs. closed access.

1. **Subject Area**: The Cancer Biology Reproducibility Project was launched in October 2013 (14) after reports from several pharmaceutical companies indicated issues in replicating published findings in cancer biology. As indicated above, the Cancer Biology Replication Project has reduced the number of studies it plans to replicate by more than 50%. Here, we compare the empirical false discovery rate of cancer journals to general medicine journals, providing a complementary measure of the replication rate.
2. **Journal Impact Factor (JIF)**: Given limited resources, most projects that attempt to replicate published studies focus on high impact papers and journals in a handful of scientific fields. However, concerns about replicability occur throughout all fields of science and levels of impact. Indeed, research published in lower impact journals may have lower rates of replicability. Here, we evaluate if JIF is associated with the empirical false discovery rate of journals.
3. **Open Access vs. closed access**: The prevalence of Open Access journals, where research is published and available to readers without a subscription or article fee, has increased considerably over the past decade (21). The number of predatory journals, which exploit the Gold Open Access model by publishing with the primary purpose of collecting submission fees to make a profit, has also increased dramatically (22, 23). While fees are common in Gold Open Access journals to remove pay walls, reputable Open Access journals have a thorough peer-review process while predatory journals have little to no peer review. Some have raised concerns that reputable Open Access journals may be letting peer-review standards fall to compete with predatory journals (23–27). Here we evaluate whether Open Access journals from InCites (28) have a higher empirical false discovery rate than journals that are not Open Access (i.e. closed access).

## Methods - Framework

Leek and Jager’s (2013) (20) method uses p-values from abstracts to arrive at an empirical false discovery rate estimate per journal per year. P-values that fall below a given significance threshold, α, are defined as positive test results and are included in the false discovery rate estimation. Within this set of positive results, results can be true or false. True discovery p-values are assumed to follow a truncated Beta distribution (tBeta) with possible observable values between 0 and *α* and with shape parameters *a* and *b*. False discoveries are assumed to follow a uniform distribution (U) between 0 and α. The true discovery and false discovery distributions are combined with mixing parameter π_0_, which is the proportion of p-values that belong to the Uniform (false discovery) distribution. If we assume that the distribution of p-values is continuous on the interval (0, 1), the combined distribution for all positive test results (i.e. p-values less than α) is:

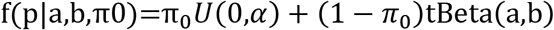

Where *a* > 0, *b* > 0 and 0 < π_0_ < 1. Using the Expectation-Maximization (EM) algorithm, the maximum likelihood estimates are simultaneously estimated for the shape parameters *a, b* and the false discovery rate, π_0_. Journal articles often do not report exact p-values (e.g. p = 0.0123); adjustments are made to the likelihood function to accommodate rounded (e.g. p = 0.01) or truncated p-values (e.g. p < 0.05). Two indicator variables are used to indicate either rounded p-values or truncated p-values. P-values that are rounded or truncated have their likelihood evaluated by integrating over all values that could possibly lead to the reported value (e.g., for p < 0.05, the associated probability is 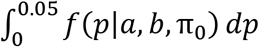; for p = 0.01, the associated probability is 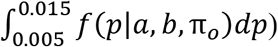. P-values are classified as rounded if the reported value has two or fewer decimal places, and as truncated if the value was read following a < or ≤ character in the text. For more details, see the Supplemental Materials of Jager and Leek (20).

## Methods - Application

We selected journals from InCites (28) using the following criteria for each journal during the years 2011-2015: available two-year JIF score, published in English, categorized as General & Internal medicine, Research & Experimental medicine, or Oncology according to InCite’s subject tags, and listed as available on the PubMed online database as of August 2017. FDR was calculated on 143 journals. The EM algorithm did not converge for one or more years for 35 journals, resulting in no FDR estimate. These journals were removed from further consideration, resulting in a final sample of 108 journals with 36,565 abstracts. InCites was used to classify journals as Open Access or closed access for each year of the study. For example, a journal marked as “Open Access since 2013” will be marked as Open Access only for the years 2013-2015. All available abstracts from 2011-2015 were collected from the online PubMed database using E-Utilities from the National Center for Biotechnology Information (29). For more details on journal selection, see Supplemental Materials.

Similar to Jager and Leek, p-values were scraped from the abstracts using regular expressions, which searched the abstract text for incidences of the phrases “p = ”, “p < ”, and “p ≤ ”. The strings following these phrases were collected and presumed to be a reported p-value. These strings were cleaned to remove excess punctuation or spacing characters and the remaining value was converted to numeric entry in scientific notation. The source code provided by Jager and Leek (20) was updated to include additional standardizing of notation and formatting in the abstracts, including scientific notation, punctuation, and spacing characters, before searching for p-values. This reduced the number of search errors from misread characters. Other than this addition, no changes were made to Jager and Leek’s original algorithm for estimating FDR. Details, including all notational substitutions, can be found in the source code available at https://github.com/laurenhall/fdr.

To identify and estimate differences in false discovery rate by journal characteristic, we applied a linear mixed effects model with the estimated false discovery rate as the outcome and a random effect by journal to account for multiple observations from each journal for each year. We fit three models: one global model with journal subject area as a covariate (1 for oncology and 0 otherwise), and two models stratified by journal subject area (oncology and medicine). Within each model, the following covariates were included: year, JIF, and Open Access status (1 if Open Access and 0 otherwise). Interaction terms between journal subject area, Open Access status, and JIF were considered for the combined model, and the interaction between Open Access status and JIF was considered for the stratified models. We then performed backwards selection on the interaction terms by assessing the significance of higher-order interaction terms first. We began with three-way and then two-way interaction terms, removing any that did not contribute significantly to the model (i.e. p-value ≤ 0.05). All main effects were left in the model regardless of significance (details and models in Supplemental). A nominal significance threshold of α = 0.05 was used to assess significance. The significance of journal subject area in the unstratified model was tested with a likelihood ratio test comparing the full model (Table 2) to a reduced model with both subject and subject by JIF removed.

To check for consistency and to ensure that our results were not driven by unusual journal characteristics, each of the three models was fit to four data sets: (1) all journals (N = 108); (2) excluding Open Access journals that were not Open Access for all five study years (N = 105); (3) excluding journals that produced an estimated false discovery rate of approximately zero (N = 97); (4) excluding both Open Access journals that were not Open Access for all five study years and journals that produced an estimated false discovery rate of approximately zero (N = 94). Models using data from (4) are shown in the Results section. Descriptions, descriptive statistics, and distributions of these four data subsets are in Supplemental Tables S1-S6 and Supplemental Figures S1-S4.

## Results

The number of journals by subject area and Open Access status included in the final model is in Table 1. A full list of journals and descriptive information is included in Supplemental Tables S7-S9.

**Table 1.** Journal Types

|  | Oncology | Medicine | Total |
| --- | --- | --- | --- |
| Open Access | 12 | 11 | 23 |
| Closed Access | 45 | 26 | 71 |
| Total | 57 | 37 | 94 |

Results of the global model are shown in Table 2. Using a likelihood ratio test to compare the global model to a reduced model with oncology and oncology by JIF removed, we find a significant difference in false discovery rate between oncology and general medicine journals (χ^2^= 24.355, df = 2, p = 5.14E-6). On average, oncology journals have a higher estimated false discovery rate compared to general medicine journals. There is also a significant interaction between subject and JIF (p = 0.028) suggesting a negative relationship between JIF and false discovery rate exists for oncology journals, but not for medicine journals. Figure 1 shows the relationship between JIF and empirical false discovery rate by journal subject area.

**Table 2.** Global Model, All Journal Types

|  | Estimate | Std. Error | T-Value | P-Value | 95% CI |
| --- | --- | --- | --- | --- | --- |
| **Year** | -0.020 | 0.004 | -0.595 | 0.552 | (-0.009, 0.005) |
| **Open Access** | 0.018 | 0.016 | 1.142 | 0.256 | (-0.014, 0.051) |
| **JIF** | -0.001 | 0.001 | -0.505 | 0.614 | (-0.002, 0.001) |
| **Oncology** | 0.092 | 0.019 | 4.992 | 1.432E-06 | (0.056, 0.129) |
| **JIF \* Onc.** | -0.005 | 0.002 | -2.208 | 0.028 | (-0.010, -0.001) |

**Fig 1.**
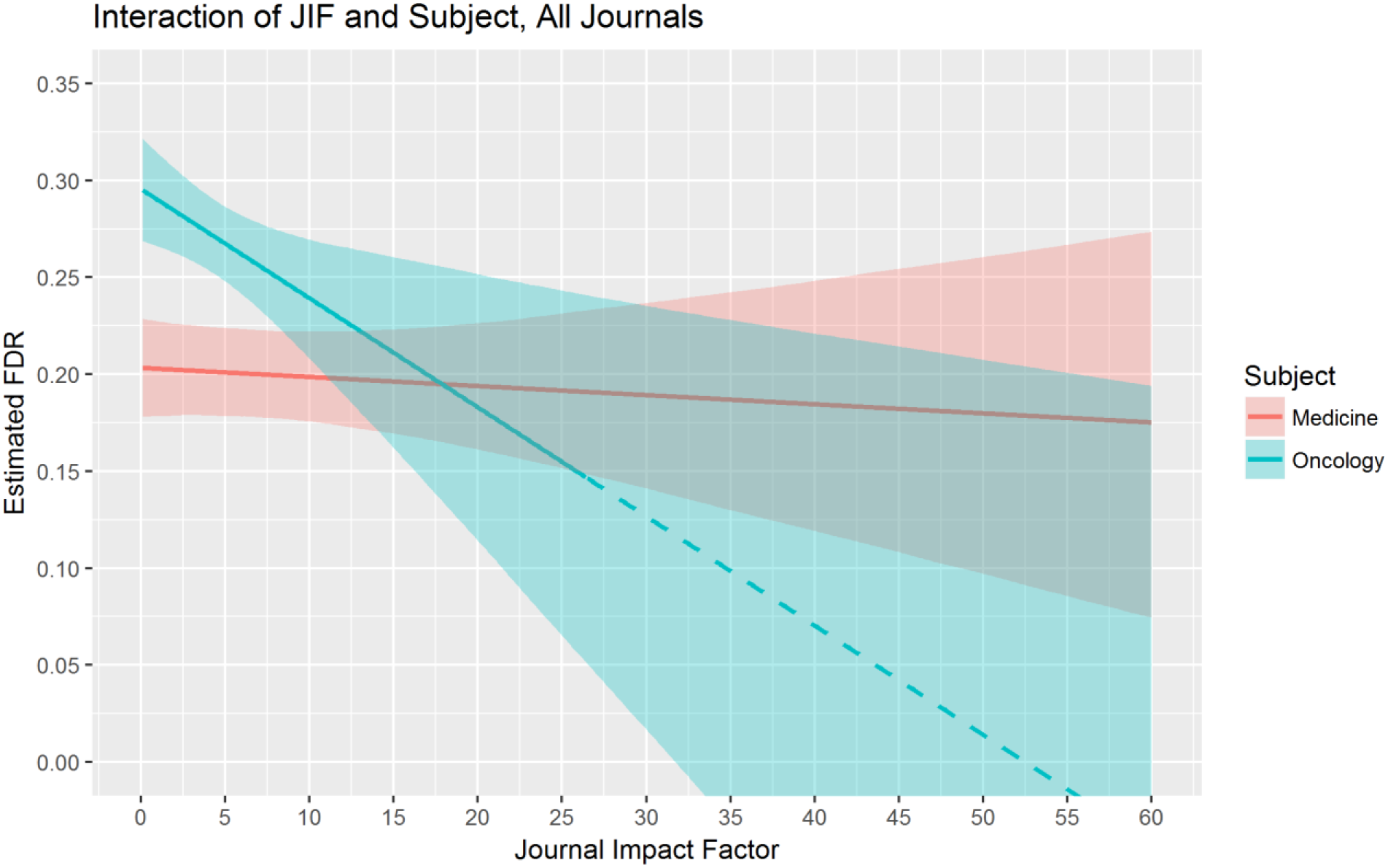
JIF and false discovery rate by subject area. Estimated linear mixed effects regression from the global model with 95% bootstrapped confidence bands. General medicine journals (red), oncology journals (blue); solid line is the predicted relationship between false discovery rate and JIF adjusting for year and open access status. The dashed blue line represents extrapolated predictions beyond the observed maximum JIF value of 26.51 for oncology journals.

Given the global model, an oncology journal with a two-year impact factor of 10 would have an estimated false discovery rate 0.042 higher than a general medicine journal with the same JIF. An oncology journal with an impact factor of 5 would have an estimated false discovery rate 0.067 higher than a comparable general medicine journal. For a journal with 1,000 reported p-values less than 0.05, this results in approximately 40 or 60 more false discovery p-values respectively compared to a general medicine journal in the same year and with the same JIF.

For the stratified model for oncology journals (Table 3), we see similar results to the global model in Table 2. We find a significant association between estimated false discovery rate and JIF with lower impact factor associated with higher false discovery rates (p = 0.012). Given this, we expect that, all else held constant, an oncology journal with a JIF of 10 would have a lower FDR of 0.03 on average compared to an oncology journal with a JIF of 5. There is no significant relationship between JIF and false discovery rate for the general medicine stratified model (Table 4, p = 0.631).

**Table 3.** Stratified: Oncology Journals

|  | Estimate | Std. Error | T-Value | P-Value | 95% CI |
| --- | --- | --- | --- | --- | --- |
| Year | -0.001 | 0.005 | -0.327 | 0.744 | (-0.011, 0.008) |
| Open Access | 0.014 | 0.023 | 0.606 | 0.547 | (-0.032, 0.059) |
| JIF | -0.006 | 0.002 | -2.548 | 0.012 | (-0.010, -0.001) |

**Table 4.**
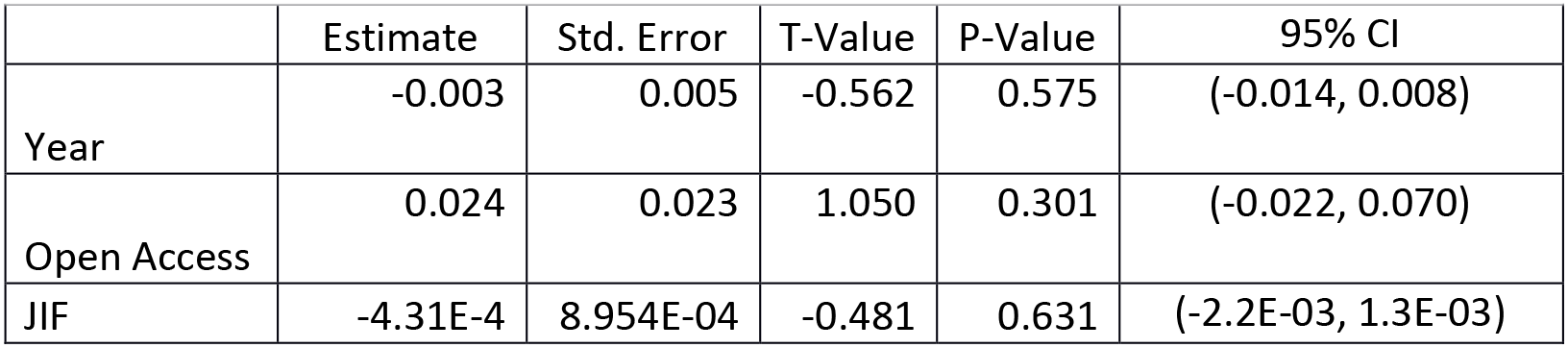
– Stratified: Medical Journals

|  | Estimate | Std. Error | T-Value | P-Value | 95% CI |
| --- | --- | --- | --- | --- | --- |
| Year | -0.003 | 0.005 | -0.562 | 0.575 | (-0.014, 0.008) |
| Open Access | 0.024 | 0.023 | 1.050 | 0.301 | (-0.022, 0.070) |
| JIF | -4.31E-4 | 8.954E-04 | -0.481 | 0.631 | (-2.2E-03, 1.3E-03) |

All secondary models, including those fit before implementing backwards selection, show results consistent with the models above and can be found in the supplemental materials (Supplemental Materials and Supplemental Tables S10-S26).

## Discussion

Using over 30,000 abstracts in 94 journals, we assessed whether journal subject area, impact factor, and Open Access status are associated with the empirical false discovery rate. We find a significantly higher empirical false discovery rate for cancer journals relative to general medicine journals and a significant inverse relationship between JIF and empirical false discovery rate within oncology journals only. Within Oncology journals, a one-unit difference in JIF is associated with an average increase in FDR of 0.006. Considering a common threshold used for the false discovery rate is 0.05, an increase of 0.006 is large. Indeed, for a ten-unit change in JIF, the average expected increase in FDR is 0.06, which would more than double an FDR of 0.05 to 0.11. These results are in line with previous reports that suggest difficulty replicating published cancer research (2). As these models assess the average relationship between factors and the empirical false discovery rate, these results do not implicate all oncology journals or journals with low JIF. Rather, these results suggest that more effort and higher standards are needed in the field of oncology research and that special attention may be needed for journals with lower impact factors.

We find no significant evidence of a relationship between Open Access status and false discovery rate. This result does not preclude the possibility that a small number of Open Access journals have a high false discovery rate. Rather this result suggests that, after adjusting for JIF, there is no significant evidence of either a systematically higher empirical false discovery rate across Open Access journals as a group or an extremely high empirical false discovery rate for a small number of Open Access journals.

There are several limitations to our study. We do not investigate patterns in the estimated false discovery rates for individual journals; rather, we assess whether certain journal characteristics (i.e. subject area, journal impact factor, Open Access status) are associated, on average, with empirical false discovery rate. Additionally, this study was performed on a sample of English-speaking journals from the field of medical research with Open Access journals from InCites for each subject area of interest. While outside of the scope of this study, increasing the sample to include non-English speaking journals, other subject areas within medicine, or repeating the study in subject areas outside of medicine would provide additional information about the relationship between the empirical false discovery rate and journal characteristics. Finally, while our inclusion of Open Access status was motivated by the increase in predatory journals, we do not directly study predatory journals here. We anticipate that our sample may underrepresent predatory journals as predatory journals are often excluded from reputable journal curation sites such as InCites. Further, restricting to English-speaking journals may exclude the majority of predatory journals that have been shown to originate in Asia and Africa (30, 31).

As Leek and Jager state in their 2017 Annual Review Stats paper (32), p-values can be presented and even manipulated in ways that can influence or call into question the accuracy of their method’s false discovery rate estimates. Here, we do not focus on the accuracy and precision of individual p-values and false discovery rates. Instead, we compare the average false discovery rate estimates by various journal characteristics. A critical assumption for our models is that any bias in the p-values is consistent between journals. It is possible, although we believe unlikely, that journal characteristics not related to the false discovery rate may change the distribution of observed p-values and thus influence the estimated false discovery rate.

We were able to complete the research presented here because Jager and Leek adhered to the highest standards of reproducible research by making their code publicly available and providing complete statistical methods. We strive to do the same here by providing complete statistical details in the supplemental section and our R code on GitHub (https://github.com/laurenhall/fdr). We hope that others will use our code and statistical details to improve upon our work and to complete research investigating patterns in the empirical false discovery rate.

Here, we investigated the relationship between the empirical false discovery rate of journals and journal subject area, JIF, and Open Access status. We find that cancer journals have a higher empirical false discovery rate compared to general medicine journals with the false discovery rate for cancer journals increasing as the JIF decreases. We do not find any significant evidence of a different empirical false discovery rate for Open Access vs. closed access journals. Given its efficiency and ability to incorporate a large and comprehensive set of published studies, the statistical framework we use here is complementary to large-scale replication studies. We hope that our approach will enable other researchers to assess the empirical false discovery rate across a wider array of disciplines and journal attributes providing insight into the patterns of replicability across science and ultimately guidance as to where more resources, higher standards, and training are needed.

## References

1. Baker M. Psychology’s reproducibility problem is exaggerated - say psychologists. Nature. 2016;267.

2. Begley CG, Ellis LM. Drug development: Raise standards for preclinical cancer research. Nature. 2012;483(7391):531–3.

3. Ioannidis JP. Why most published research findings are false. PLoS Med. 2005;2(8):e124.

4. Ioannidis JP. Contradicted and initially stronger effects in highly cited clinical research. JAMA. 2005;294(2):218–28.

5. Ioannidis JP, Allison DB, Ball CA, Coulibaly I, Cui X, Culhane AC, et al. Repeatability of published microarray gene expression analyses. Nat Genet. 2009;41(2):149–55.

6. Open Science Collaboration. PSYCHOLOGY. Estimating the reproducibility of psychological science. Science. 2015;349(6251):aac4716.

7. Pusztai L, Hatzis C, Andre F. Reproducibility of research and preclinical validation: problems and solutions. Nat Rev Clin Oncol. 2013;10(12):720–4.

8. Baggerly K, Coombes K. Deriving chemosensitivity from cell lines: Forensic bioinformatics and reproducible research in high-throughput biology. The Annals of Applied Statistics. 2009:1309–34.

9. National Academy of Sciences. Reproducibility and Replicability in Science 2018 [Available from: http://sites.nationalacademies.org/dbasse/bbcss/reproducibilityandreplicabilityinscience/index.htm.

10. Mesirov JP. Computer science. Accessible reproducible research. Science. 2010;327(5964):415–6.

11. Peng RD. Reproducible research in computational science. Science. 2011;334(6060):1226–7.

12. Peng RD, Dominici F, Zeger SL. Reproducible epidemiologic research. Am J Epidemiol. 2006;163(9):783–9.

13. Benjamin DJ, et al. Redefine statistical significance. Nature Human Behavior. 2018;2(1):6.

14. eLife Sciences Publications. Reproducibility Project: Cancer Biology 2018 [Available from: https://elifesciences.org/collections/9b1e83d1/reproducibility-proiect-cancer-biology.

15. Camerer CF, et al. Evaluating replicability of laboratory experiments in economics. Science. 2016;351(6280):1433–6.

16. The Social Sciences Replication Project. Social Sciences Replication Project 2016 [Available from: http://www.socialsciencesreplicationproiect.com/.

17. Camerer CF, et al. Evaluating the replicability of social science experiments in Nature and Science between 2010 and 2015. Nature Human Behavior. 2018:1.

18. Patil P, Peng RD, Leek JT. What Should Researchers Expect When They Replicate Studies? A Statistical View of Replicability in Psychological Science. Perspect Psychol Sci. 2016;11(4):539–44.

19. Kaiser J. Plan to replicate 50 high-impact cancer papers shrinks to just 18. Science magazine. 2018 July 31, 2018.

20. Jager LR, Leek JT. An estimate of the science-wise false discovery rate and application to the top medical literature. Biostatistics. 2013;15(1):1–12.

21. Poltronieri E, Bravo E, Curti M, Ferri M, Mancini C. Open access publishing trend analysis: statistics beyond the perception. Information research. 2016;21(2).

22. Haug C. The downside of open-access publishing. N Engl J Med. 2013;368(9):791–3.

23. Shen C, Björk BC. “Predatory” open access: A longitudinal study of article volumes and market characteristics. BMC Medicine. 2015;13(1):1–15.

24. Beall J. What I learned from predatory publishers. Biochem Med (Zagreb). 2017;27(2):273–8.

25. Beall J. Predatory publishers are corrupting open access. Nature. 2012;489(7415):179.

26. Singh Chawla D. The undercover academic keeping tabs on ‘predatory’ publishing. Nature. 2018;555(7697):422–3.

27. Swauger S. Open access, power, and privilege: A response to “What I learned from predatory publishing.”. College & Research Libraries News. 2017;78(11):603.

28. Thompson-Reuters. InCites 2018 [Available from:http://incites.thomsonreuters.com.

29. National Center for Biotechnological Information. Entrez Programming Utilities 2010 [Available from:https://www.ncbi.nlm.nih.gov/books/NBK25501/.

30. Hansoti BaL M., and Murphy, L. Discriminating between legitimate and predatory open access journals: Report from the International Federation for Emergency Medical Research. Western Journal of Emergency Medicine. 2016;17(5):497–507.

31. Omobowale A, Akanle A, Adeniran I, Olayinka K. Peripheral scholarship and the context of foreign paid publishing in Nigeria. Current Sociology. 2014;62:666–84.

32. Leek JT, Jager LR. Is Most Published Research Really False? Annu Rev Stat Appl. 2017;4:109–22.

