## Supplemental Materials for "Analysis of >30,000 abstracts suggests higher false discovery rates for oncology journals, especially those with low impact factors" (from another version of this preprint: version 2, 2019-01-03)

**Supplementary Materials**

**Detailed journal selection criteria**

One hundred and sixty-six journals were identified by accessing the Thompson-Reuters InCites journal list of journals that had a subject marked as “Oncology”, “Medicine – General & Internal”, or “Medicine – Research & Experimental”. The full list of journals is in the supplementary file “Journals.csv” (<https://github.com/laurenhall/fdr>) and was compiled on August 23, 2017.

The abstracts for these journals were scraped from PubMed for the years 2011-2015. Journals that had no abstracts available for one of more of these years were eliminated (13 journals): *Advancement in Clinical and Experimental Medicine, Analytic Cell Pathology, Canadian Medical Association Journal, Cancer Immunology Research, Cancer Prevention Research, Cellular Oncology, Clinical Medicine, Concoimmunology, Experimental Biological Medicine, Gynaecological Oncology, Journal of Diabetes Research, Molecular Therapy – Nucleic Acids,* and *Sexual Medicine*.

We estimated the false discovery rate for the remaining 153 journals for each of the five research years. Some journals, due to either a small number of p-values or not enough unique observations (as can occur when most reported p-values are truncated), did not produce false discovery estimates for one or more years. These journals were eliminated (35 journals): *Cancer Cell, Cancer Cell International, Cancer Discovery, Cancer Imaging, Cancer Journal, Clinical Journal of Oncological Nursing, Current Molecular Medicine, Discovery Medicine, EMBO Molecular Medicine, European Journal of General Practice, Experimental Cell Research, Family Medicine, Future Oncology, Journal of Cancer Education, Journal of Clinical Investigation, Journal of Experimental Medicine, Journal of Genetic Medicine, Journal of Immunotherapy, Journal of International Medical Research, Journal of the National Comprehensive Cancer Network, Journal of the National Medical Association, Journal of the Royal Society of Medicine, MAbs, Molecular Aspects in Medicine, Nature Medicine, Oncology Research, Pigment Cell Melanoma Research, Primary Care, Seminars in Cancer Biology, Seminars in Oncology, Seminars in Radiation and Oncology, Statistics in Medicine, Stem Cell Research and Therapy, Theranostics,* and *Trends in Molecular Medicine.*

In addition to the above journals, several journals were removed from consideration due to not having a two-year journal impact factor (JIF) score available for one or more of the study years (10 journals): *American Journal of Cancer Research, Blood Cancer Journal, BMJ Open, Cancer Research and Treatment, Clinical Epigenetics, Clinics, Hormones and Cancer, International Journal of Clinical and Experimental Medicine, Journal of Cancer,* and *Journal of Ophthalmology.*

The remaining 108 journals were included in the study. The full list of journals, including estimated false discovery rate, number of abstracts received, and number of p-values identified in abstracts can be found in Table S1. Table S2 below describes the distribution of journals present in this data set by subject and Open Access status. Journals which were Open Access in some years and not others count towards both categories, proportional to how many years they spent in each state.

Table S2 – Number of Journals by Subject and Open Access status

|  | Oncology | Medicine | Total |
| --- | --- | --- | --- |
| Open Access | 12 | 11 | 23 |
| Closed Access | 49 | 30 | 79 |
| Both | 2 | 4 | 6 |
| Total | 63 | 45 | 108 |

**Reduced data sets**

In addition to the full data set consisting of 108 journals described in the previous section (i.e. dataset 1), three reduced data sets were used in this study. These data sets have one or both of the following groups of journals removed:

2) Remove journals with FDR estimates close to zero: As stated in the methods, the EM-algorithm is estimating the proportion of false discovery p-values is limited to between 0 and 1 and cannot equal 0 or 1. Estimates of the number of false discovery p-values close to zero may instead be zero. An estimated false discovery rate close to zero implies that no p-values in the sample were false discoveries. We removed journals if the expected number of false discovery p-values for a journal in a year was less than 0.01. The expected number of false discovery p-values was calculated by multiplying the total number of p-values times the estimated FDR.

Journals excluded from this data set: *Prostate Cancer and Prostatic Disease, Psycho-oncology, Radiation Oncology, Radiology and Oncology, Radiotherapy Oncology, Surgical Oncology, Target Oncology, Thoracic Cancer, Translational Oncology, Trials,* and *World Journal of Surgical Oncology*.

The full list of journals, including estimated false discovery rate, number of abstracts received, and number of p-values identified in abstracts for this reduced data set can be found in Table S3. Table S4 below describes the distribution of journals present in this data set by subject and Open Access status.

Table S4 – Number of Journals by Subject and Open Access status after removing journals with FDR close to zero

|  | Oncology | Medicine | Total |
| --- | --- | --- | --- |
| Open Access | 12 | 11 | 23 |
| Closed Access | 45 | 26 | 71 |
| Both | 1 | 2 | 3 |
| Total | 58 | 39 | 97 |

3) Remove journals that were not consistently Open Access or closed access during the study years: Some journals switched to an Open Access model during the study years. In the unfiltered data set, these journals are encoded with a 1 for Open Access for the years they were Open Access, and 0 otherwise. For this data subset, we removed these journals.

Journals excluded from this data set: *Surgical Oncology, Target Oncology, Thoracic Cancer, Translational Oncology, Trials,* and *World Journal of Surgical Oncology*.

The full list of journals, including estimated false discovery rate, number of abstracts received, and number of p-values identified in abstracts for this reduced data set can be found in Table S5. Table S6 below describes the distribution of journals present in this data set by subject and Open Access status.

Table S6 – Number of Journals by Subject and Open Access status after removing journals that switched Open Access status

|  | Oncology | Medicine | Total |
| --- | --- | --- | --- |
| Open Access | 12 | 11 | 23 |
| Closed Access | 49 | 30 | 79 |
| Total | 61 | 41 | 102 |

4) Reduced data set excluding both journals from 2 and 3: This data set excludes both sets of journals described above and is the primary dataset presented in the paper. The full list of journals, estimated false discovery rate, number of abstracts, and number of p-values can be found in Table S7. Table S8 below describes the distribution of journals present in this dat set by subject and Open Access status.

Table S8 – Number of Journals by Subject and Open Access status after removing journals that switched Open Access status and had FDR estimates close to zero

|  | Oncology | Medicine | Total |
| --- | --- | --- | --- |
| Open Access | 12 | 11 | 23 |
| Closed Access | 45 | 26 | 71 |
| Total | 57 | 37 | 94 |

Additionally, a full list of all considered journals organized by subject and group membership status can be found in Table S9.

**Model Selection and Models for Additional Data Sets**

As described in the main text, we arrived at our final model through a modified selection method. Beginning with a full model including all two-way and three-way (where appropriate) interaction terms, we removed any interactions that were not significant beginning with the three-way terms and then the two-way terms, beginning with the least significant. No main effects were removed from any of the models. The tables below contain the stages of model selection not shown in the main text, as well as models for the data sets described in the previous section to demonstrate consistency of results.

**Unstratified Model – Subject as a Covariate**

**Model Selection Details – Main Text Models**

Table S10 – Full Model, Journals with Zero Estimates and Inconsistent Open Access Status Removed

|  | **Estimate** | **Std. Error** | **T-Value** | **P-Value** | **95% CI** |
| --- | --- | --- | --- | --- | --- |
| **Intercept** | 4.821 | 7.292 | 0.661 | 0.509 | (-9.518, 19.160) |
| **Year** | -0.002 | 0.004 | -0.635 | 0.526 | (-0.009, 0.005) |
| **Open Access** | 0.058 | 0.033 | 1.781 | 0.078 | (-0.007, 0.124) |
| **JIF** | -1.641E-04 | 9.587E-04 | -0.171 | 0.864 | (-0.002, 0.002) |
| **Oncology** | 0.010 | 0.022 | 4.594 | 7.045E-06 | (0.057, 0.143) |
| **OA * JIF** | -0.008 | 0.005 | -1.546 | 0.123 | (-0.019, 0.002) |
| **OA * Onc.** | -0.073 | 0.055 | -1.336 | 0.185 | (-0.181, 0.036) |
| **JIF * Onc.** | -0.006 | 0.002 | -2.410 | 0.017 | (-0.012, -0.001) |
| **OA * JIF * Onc.** | 0.017 | 0.013 | 1.324 | 0.186 | (-0.008, 0.042) |

Table S11 – Journals with Zero Estimates and Inconsistent Open Access Status Removed, Excluding 3-Way Interaction

|  | **Estimate** | **Std. Error** | **T-Value** | **P-Value** | **95% CI** |
| --- | --- | --- | --- | --- | --- |
| **Intercept** | 4.506 | 7.295 | 0.618 | 0.537 | (-9.839, 18.851) |
| **Year** | -0.002 | 0.004 | -0.592 | 0.555 | (-0.009, 0.005) |
| **Open Access** | 0.047 | 0.031 | 1.467 | 0.146 | (-0.016, 0.109) |
| **JIF** | -2.601E-04 | 9.562E-04 | -0.272 | 0.786 | (-0.002, 0.002) |
| **Oncology** | 0.096 | 0.022 | 4.464 | 1.161E-05 | (0.053, 0.139) |
| **OA * JIF** | -0.005 | 0.005 | -1.094 | 0.2746 | (-0.015, 0.004) |
| **OA * Onc.** | -0.015 | 0.033 | -0.462 | 0.645 | (-0.081, 0.501) |
| **JIF * Onc.** | -0.005 | 0.002 | -2.199 | 0.029 | (-9.965E-03, -5.562E-04) |

Table S12 – Journals with Zero Estimates and Inconsistent Open Access Status Removed, Excluding OA * Oncology Interaction

|  | **Estimate** | **Std. Error** | **T-Value** | **P-Value** | **95% CI** |
| --- | --- | --- | --- | --- | --- |
| **Intercept** | 4.521 | 7.295 | 0.620 | 0.536 | (-9.822, 18.865) |
| **Year** | -0.002 | 0.004 | -0.593 | 0.553 | (-0.009, 0.005) |
| **Open Access** | 0.037 | 0.024 | 1.539 | 0.127 | (-0.011, 0.084) |
| **JIF** | -3.252E-04 | 9.417E-04 | -0.345 | 0.730 | (-0.002, 0.002) |
| **Oncology** | 0.091 | 0.019 | 4.910 | 1.997E-06 | (0.054, 0.128) |
| **OA * JIF** | -0.005 | 0.005 | -1.043 | 0.298 | (-0.014, 0.004) |
| **JIF * Onc.** | -0.005 | 0.002 | -2.164 | 0.031 | (-9.680E-03, -4.624E-04) |

Table S13 – Final Model, Journals with Zero Estimates and Inconsistent Open Access Status Removed, Excluding OA * JIF Interaction

|  | **Estimate** | **Std. Error** | **T-Value** | **P-Value** | **95% CI** |
| --- | --- | --- | --- | --- | --- |
| **Intercept** | 4.536 | 7.299 | 0.621 | 0.535 | (-9.816, 18.888) |
| **Year** | -0.002 | 0.004 | -0.595 | 0.552 | (-0.009, 0.005) |
| **Open Access** | -0.018 | 0.016 | 1.142 | 0.256 | (-0.014, 0.051) |
| **JIF** | -4.692E-04 | 9.296E-04 | -5.047 | 0.614 | (-0.002, 0.001) |
| **Oncology** | 0.092 | 0.012 | 4.992 | 1.432E-06 | (0.056, 0.129) |
| **JIF * Onc.** | -0.005 | 0.002 | -2.208 | 0.028 | (-9.758E-03, -5.643E-04) |

**Final Combined Models – Secondary Data Sets**

Data Set 1: All Journals

Table S14 – Final Model, All Journals

|  | **Estimate** | **Std. Error** | **T-Value** | **P-Value** | **95% CI** |
| --- | --- | --- | --- | --- | --- |
| **Intercept** | -5.301 | 7.236 | -0.733 | 0.464 | (-19.523, 8.921) |
| **Year** | 0.003 | 0.004 | 0.759 | 0.448 | (-0.004, 0.010) |
| **Open Access** | 0.222 | 0.015 | 1.465 | 0.144 | (-0.008, 0.052) |
| **JIF** | -5.215E-04 | 9.355E-04 | -0.557 | 0.578 | (-0.002, 0.001) |
| **Oncology** | 0.098 | 0.018 | 5.610 | 8.146E-08 | (0.064, 0.133) |
| **JIF * Onc.** | -0.005 | 0.002 | -2.121 | 0.035 | (-9.586E-03, -3.648E-04) |

Data Set 2: Journals with Zero Estimates removed

Table S15 – Final Model, Zero Estimates Removed

|  | **Estimate** | **Std. Error** | **T-Value** | **P-Value** | **95% CI** |
| --- | --- | --- | --- | --- | --- |
| **Intercept** | 3.024 | 7.154 | 0.423 | 0.672 | (-11.044, 17.091) |
| **Year** | -0.001 | 0.004 | -0.395 | 0.693 | (-0.008, 0.006) |
| **Open Access** | 0.022 | 0.016 | 1.381 | 0.168 | (-0.009, 0.053) |
| **JIF** | -5.736E-04 | 9.393E-04 | -0.611 | 0.542 | (-0.002, 0.001) |
| **Oncology** | 0.092 | 0.018 | 5.006 | 1.279E-06 | (0.056, 0.129) |
| **JIF * Onc.** | -0.005 | 0.002 | -2.162 | 0.031 | (-9.786E-03, -4.636E-04) |

Data Set 3: Journals with Inconsistent Open Access Status removed

Table S16 – Final Model, Inconsistent Open Access Status Removed

|  | **Estimate** | **Std. Error** | **T-Value** | **P-Value** | **95% CI** |
| --- | --- | --- | --- | --- | --- |
| **Intercept** | -5.301 | 7.236 | -0.733 | 0.464 | (-19.523, 8.921) |
| **Year** | 0.003 | 0.004 | 0.759 | 0.448 | (-0.004, 0.010) |
| **Open Access** | 0.222 | 0.015 | 1.465 | 0.144 | (-0.008, 0.052) |
| **JIF** | -5.215E-04 | 9.355E-04 | -0.557 | 0.578 | (-0.002, 0.001) |
| **Oncology** | 0.098 | 0.018 | 5.610 | 8.146E-08 | (0.064, 0.133) |
| **JIF * Onc.** | -0.005 | 0.002 | -2.121 | 0.035 | (-9.586E-03, -3.648E-04) |

**Stratified Models – Oncology**

**Model Selection Details – Main Text Models**

Table S17 – Oncology Full Model, Journals with Zero Estimates and Inconsistent Open Access Status Removed

|  | **Estimate** | **Std. Error** | **T-Value** | **P-Value** | **95% CI** |
| --- | --- | --- | --- | --- | --- |
| **Intercept** | 3.778 | 9.827 | 0.384 | 0.701 | (-15.588, 23.144) |
| **Year** | -0.002 | 0.005 | -0.355 | 0.723 | (-0.011, 0.008) |
| **Open Access** | -0.014 | 0.045 | -0.317 | 0.752 | (-0.105, 0.076) |
| **JIF** | -0.006 | 0.002 | -2.620 | 0.009 | (-0.011, -0.002) |
| **OA * JIF** | 0.009 | 0.012 | 0.716 | 0.474 | (-0.015, 0.032) |

Table S18 – Oncology Final Model, Journals with Zero Estimates and Inconsistent Open Access Status Removed, excluding OA * JIF interaction

|  | **Estimate** | **Std. Error** | **T-Value** | **P-Value** | **95% CI** |
| --- | --- | --- | --- | --- | --- |
| **Intercept** | 3.498 | 9.822 | 0.356 | 0.722 | (-15.855, 22.852) |
| **Year** | -0.002 | 0.005 | -0.327 | 0.744 | (-0.011, 0.008) |
| **Open Access** | -0.014 | 0.023 | 0.686 | 0.547 | (-0.032, 0.059) |
| **JIF** | -0.006 | 0.002 | -2.548 | 0.012 | (-0.010, -0.001) |

**Final Stratified Models – Oncology – Secondary Data Sets**

Data Set 1: All Journals

Table S19 – Oncology Final Model, All Journals

|  | **Estimate** | **Std. Error** | **T-Value** | **P-Value** | **95% CI** |
| --- | --- | --- | --- | --- | --- |
| **Intercept** | -7.656 | 9.997 | -0.766 | 0.445 | (-27.346, 12.034) |
| **Year** | 0.004 | 0.005 | 0.795 | 0.428 | (-0.006, 0.014) |
| **Open Access** | 0.019 | 0.022 | 0.871 | 0.385 | (-0.024, 0.061) |
| **JIF** | -0.006 | 0.002 | -2.547 | 0.012 | (-0.010, -0.001) |

Data Set 2: Journals with Zero Estimates removed

Table S20 – Oncology Final Model, Zero Estimates Removed

|  | **Estimate** | **Std. Error** | **T-Value** | **P-Value** | **95% CI** |
| --- | --- | --- | --- | --- | --- |
| **Intercept** | 2.596 | 9.729 | 0.227 | 0.790 | (-16.574, 21.766) |
| **Year** | -0.001 | 0.005 | -0.237 | 0.813 | (-0.011, 0.008) |
| **Open Access** | 0.017 | 0.022 | 0.738 | 0.461 | (-0.028, 0.061) |
| **JIF** | -0.006 | 0.002 | -2.534 | 0.012 | (-0.010, -0.001) |

Data Set 3: Journals with Inconsistent Open Access Status removed

Table S21 – Oncology Final Model, Inconsistent Open Access Status Removed

|  | **Estimate** | **Std. Error** | **T-Value** | **P-Value** | **95% CI** |
| --- | --- | --- | --- | --- | --- |
| **Intercept** | -4.456 | 10.097 | -0.441 | 0.659 | (-24.345, 15.433) |
| **Year** | -0.002 | 0.005 | 0.470 | 0.639 | (-0.008, 0.012) |
| **Open Access** | 0.014 | 0.022 | 0.639 | 0.525 | (-0.030, 0.058) |
| **JIF** | -0.005 | 0.002 | 02.472 | 0.014 | (-0.010, -0.001) |

**Stratified Models – Medicine**

**Model Selection Details – Main Text Models**

Table S22 – Medicine Full Model, Journals with Zero Estimates and Inconsistent Open Access Status Removed

|  | **Estimate** | **Std. Error** | **T-Value** | **P-Value** | **95% CI** |
| --- | --- | --- | --- | --- | --- |
| **Intercept** | 6.565 | 10.718 | 0.613 | 0.541 | (-14.618, 27.749) |
| **Year** | -0.003 | 0.005 | -0.595 | 0.553 | (-0.014, 0.007) |
| **Open Access** | -0.588 | 0.031 | 1.902 | 0.065 | (-0.004, 0.122) |
| **JIF** | -1.588E-04 | 9.040E-04 | -0.176 | 0.861 | (-0.002, 0.002) |
| **OA * JIF** | -0.008 | 0.005 | -1.659 | 0.099 | (-0.018, 0.002) |

Table S23 – Medicine Final Model, Journals with Zero Estimates and Inconsistent Open Access Status Removed, excluding OA * JIF interaction

|  | **Estimate** | **Std. Error** | **T-Value** | **P-Value** | **95% CI** |
| --- | --- | --- | --- | --- | --- |
| **Intercept** | 6.239 | 10.762 | 0.580 | 0.563 | (-15.030, 27.509) |
| **Year** | -0.003 | 0.005 | -0.562 | 0.575 | (-0.014, 0.008) |
| **Open Access** | 0.024 | 0.023 | 1.050 | 0.301 | (-0.022, 0.070) |
| **JIF** | -4.301E-04 | 8.954E-04 | -0.481 | 0.631 | (-0.002, 0.001) |

**Final Stratified Models – Medicine – Secondary Data Sets**

Data Set 1: All Journals

Table S24 – Medicine Final Model, All Journals

|  | **Estimate** | **Std. Error** | **T-Value** | **P-Value** | **95% CI** |
| --- | --- | --- | --- | --- | --- |
| **Intercept** | -1.875 | 10.316 | -0.182 | 0.856 | (-22.232, 18.483) |
| **Year** | 0.001 | 0.005 | 0.200 | 0.842 | (-0.009, 0.011) |
| **Open Access** | 0.025 | 0.021 | 1.187 | 0.237 | (-0.017, 0.0671) |
| **JIF** | -5.100E-04 | 9.279E-04 | -0.550 | 0.583 | (-0.002, 0.001) |

Data Set 2: Journals with Zero Estimates removed

Table S25 – Medicine Final Model, Zero Estimates Removed

|  | **Estimate** | **Std. Error** | **T-Value** | **P-Value** | **95% CI** |
| --- | --- | --- | --- | --- | --- |
| **Intercept** | 3.844 | 10.398 | 0.370 | 0.712 | (-16.698, 24.386) |
| **Year** | -0.002 | 0.005 | -0.035 | 0.726 | (-0.012, 0.008) |
| **Open Access** | 0.028 | 0.022 | 1.283 | 0.201 | (-0.015, 0.072) |
| **JIF** | -5.338E-04 | 8.997E-04 | -0.593 | 0.554 | (-0.002, 0.001) |

Data Set 3: Journals with Inconsistent Open Access Status removed

Table S26 – Medicine Final Model, Inconsistent Open Access Status Removed

|  | **Estimate** | **Std. Error** | **T-Value** | **P-Value** | **95% CI** |
| --- | --- | --- | --- | --- | --- |
| **Intercept** | -1.854 | 10.494 | -0.177 | 0.860 | (-22.586, 18.878) |
| **Year** | 0.001 | 0.005 | 0.194 | 0.846 | (-0.009, 0.011) |
| **Open Access** | 0.032 | 0.024 | 1.349 | 0.185 | (-0.016, 0.080) |
| **JIF** | 3.310E-04 | 9.486E-04 | -0.349 | 0.728 | (-0.002, 0.002) |

**Supplemental Figures**

**Distribution of Scraped P-Values**

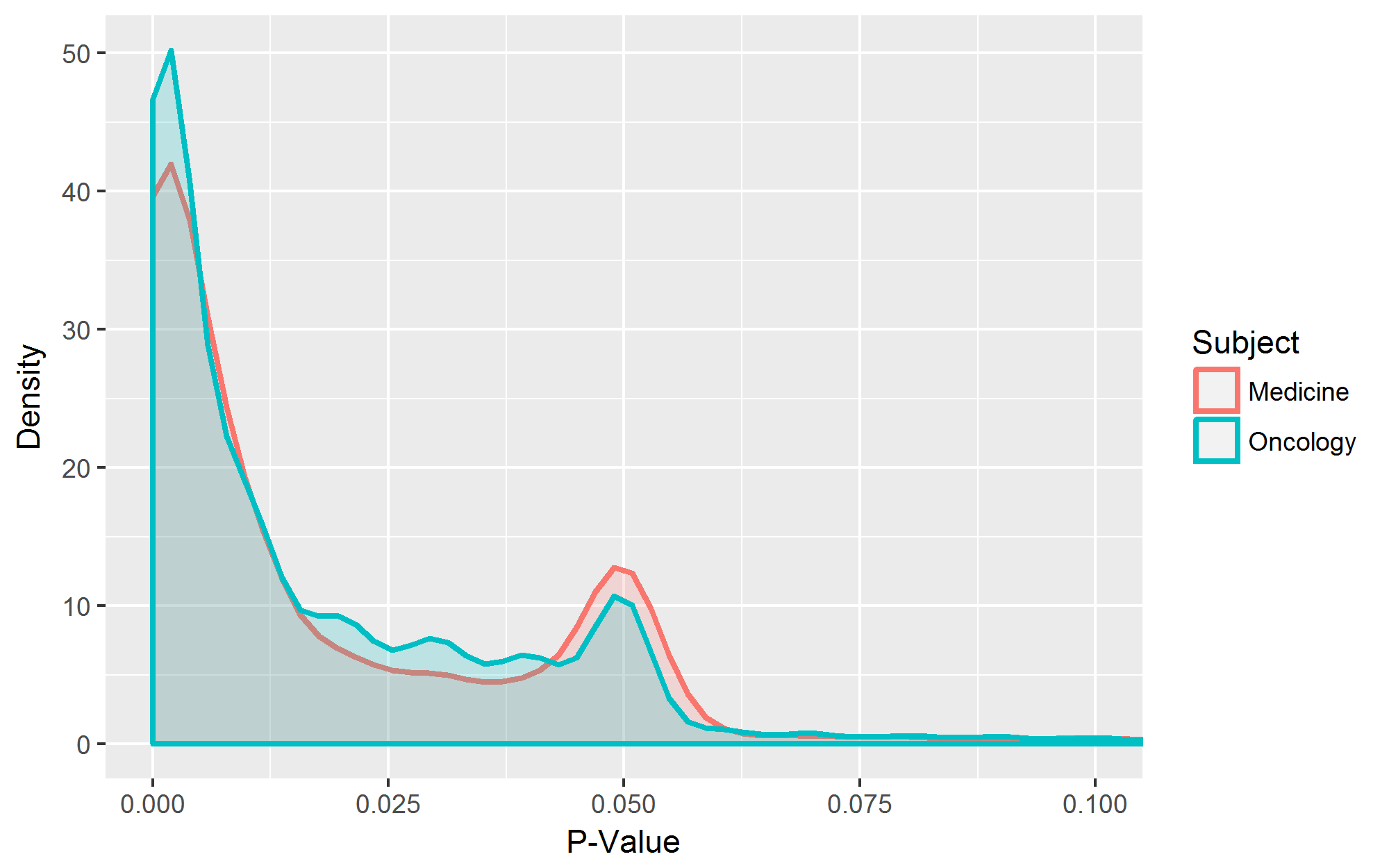

**Fig S1 – Distribution of scraped p-values between 0 and 0.1 by Subject.** All journals and all years included. Medicine journals (red) and Oncology journals (blue). The peaks at 0.001 and 0.05 represent large numbers of p-values reported as P < 0.001 or P < 0.05, respectively.

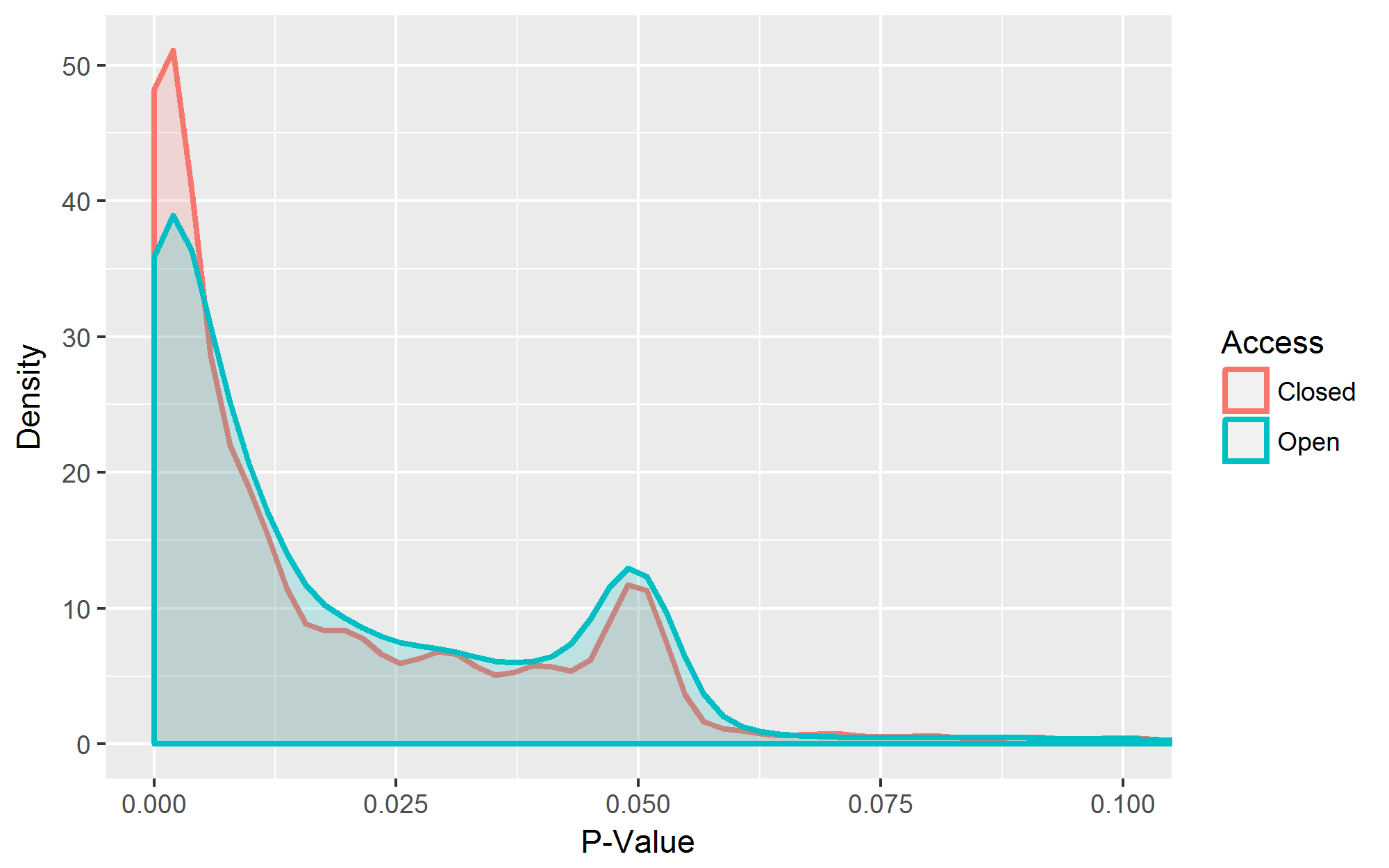

**Fig S2 – Distribution of scraped p-values between 0 and 0.1 by Access Type.** Journals with “Both” access type excluded. Closed access (red) vs. Open Access (blue). The peaks at 0.001 and 0.05 represent large numbers of p-values reported as P < 0.001 or P < 0.05, respectively.

**Distribution of FDR Estimates by Journal Impact Factor**

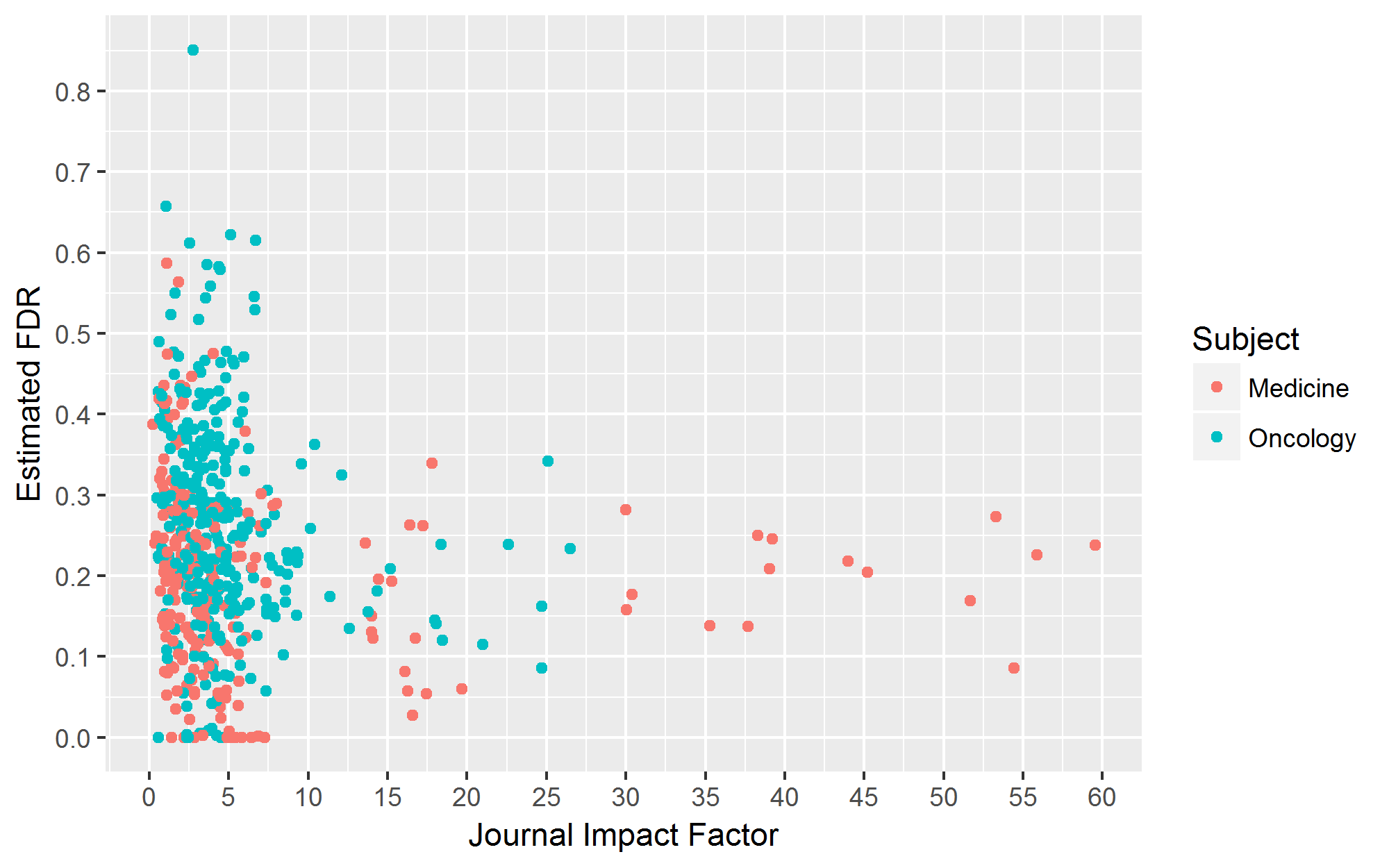

**Fig S3 – Estimated FDR and JIF by Subject.** All journals and all years included. Medicine journals (red) and Oncology journals (blue). JIF for medicine journals ranged from 0.206 to 56.558. JIF for oncology journals ranged from 0.474 to 26.509.

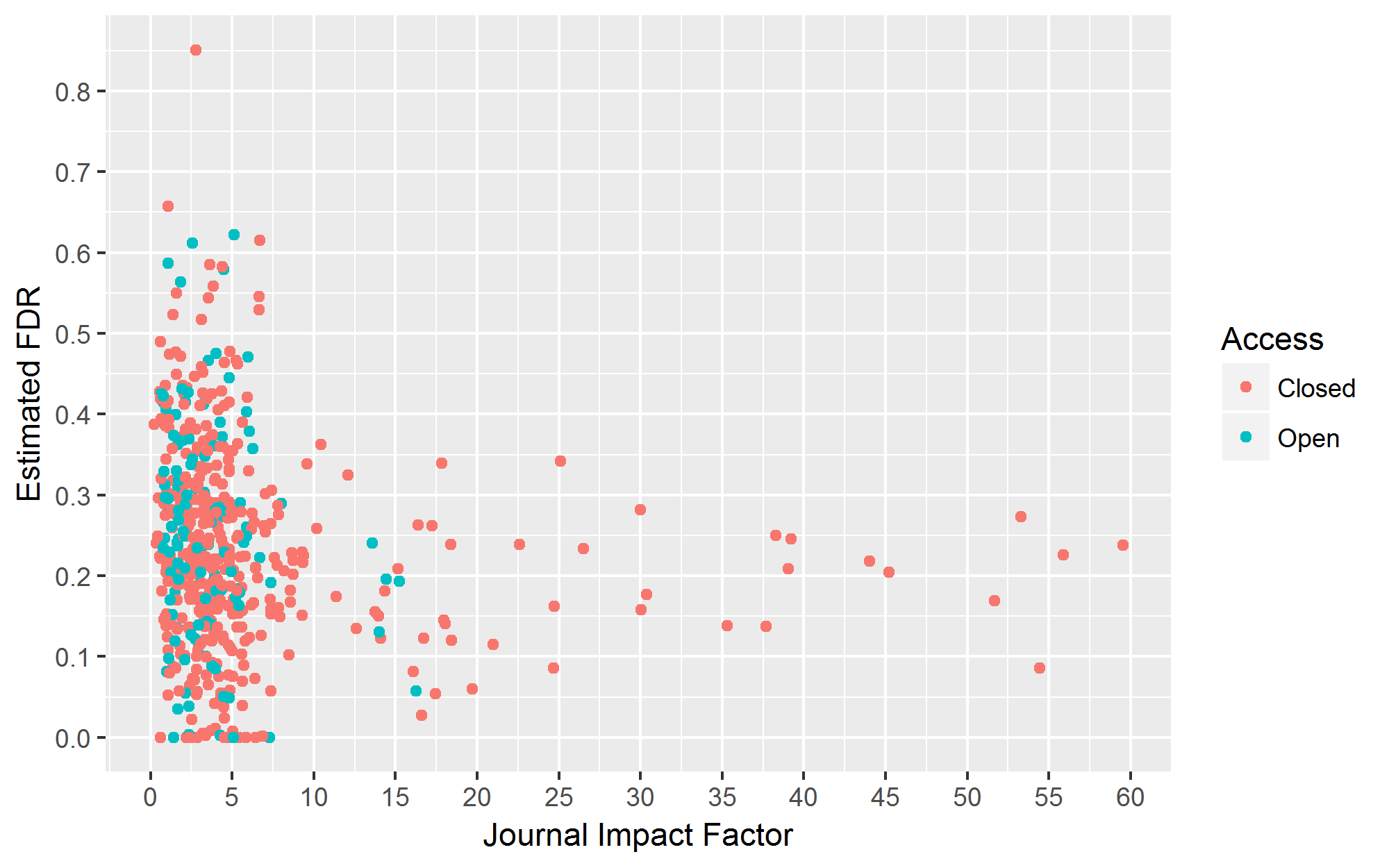

**Fig S4 – Estimated FDR and JIF by Access Type**. Journals with “Both” access type excluded. Closed access (red) vs. Open Access (blue). JIF for closed access journals ranged from 0.206 to 56.558. JIF for Open Access journals ranged from 0.656 to 16.269.
